# Spatial differentiation of background matching strategies along a Late Pleistocene range expansion route

**DOI:** 10.1101/2022.05.27.493691

**Authors:** Giada Spadavecchia, Andrea Chiocchio, David Costantini, Anita Liparoto, Roberta Bisconti, Daniele Canestrelli

## Abstract

Late Pleistocene climate changes have deeply impacted the range dynamics of temperate species. While the genetic legacy of these dynamics has been widely investigated, little is known about their phenotypic consequences. Anti-predatory strategies offer intriguing opportunities to study phenotypic evolution in response to dispersal dynamics since the ability to avoid predation can be pivotal for populations colonizing new environments. Here we investigated the spatial differentiation of background colour matching strategies along a Late Pleistocene range expansion route of a temperate species, the Tyrrhenian tree frog *Hyla sarda*. Using common-garden experiments, we investigated whether individuals sampled in the source area (Sardinia) and individuals sampled in the newly founded area (Corsica) differ in two components of the camouflage strategy: colour change abilities and background choice behaviour. We found a remarkable spatial structure in both colour change abilities and background choice behaviour, across the expansion range. Tree frogs from the source area displayed higher colour change abilities and a more pronounced preference for a greener background, with respect to tree frogs from the newly colonized area. We discuss these results in the context of the spatial and demographic components of the expansion dynamics. Our results support the intriguing hypothesis that Late Pleistocene biogeographic history might be an overlooked major player in shaping current spatial patterns of phenotypic traits variation across animal populations.

## 1. Introduction

Multiple evidence from biogeographic and eco-evolutionary research documented that virtually all species responded to Pleistocene climatic changes with marked oscillations of their geographic distribution (Taberlet et al., 1998; Hewitt, 2004a; Schmitt, 2007; Fløjgaard et al., 2009; Hofreiter & Stewart, 2009). During periods of unsuitable climatic conditions, species’ ranges contracted into climatic refugia, whereas range expansions followed the re-establishment of suitable conditions (Hewitt, 1996; 2000; 2011a; Weiss & Ferrand, 2007). The legacies of range dynamics on species genetic structures have been widely investigated during the last decades (Hewitt, 2000; Avise, 2009; Hewitt, 2011b; Lucati et al., 2020; Chiocchio et al., 2021), whereas evidence on their role in moulding intra-specific phenotypic variation is scarce. Yet, phenotypic traits are likely to vary in response to dispersal processes, especially during range expansions (Canestrelli et al., 2016a). Indeed, expanding populations experience different selection regimes along the expansion route, and substantial changes in their phenotypic makeup can occur by spatial sorting of traits implicated in density-dependent processes, such as dispersal-related traits, traits involved in sexual selection (including behavioural traits), and anti-predatory strategies (Cobben et al., 2015; Canestrelli et al., 2016b; Szűcs et al., 2017; Miller et al., 2020).

Background matching is an anti-predatory strategy consisting of body colours or patterns that resemble those in the surrounding environment, decreasing the probability of being detected by predators (Merilaita & Stevens, 2011; Merilaita et al., 2017; Ruxton, 2019). In maThus, analysing spatial patterns of intenry instances, background matching strategies have a clear behavioural component, since animals actively seek an appropriate background resembling their body colouration (Kjernsmo et al., 2012; Marshall et al., 2016; Stevens et al., 2017; Stevens & Ruxton. 2019). Alternatively, chromatic resemblance to specific substrates can be achieved through physiologically mediated colour change abilities (Duarte et al., 2017; Stevens, 2016; Sköld et al., 2013; Stuart-Fox& Moussali, 2011). Background choice and colour change are often combined, especially in animals with limited body pattern repertoire or low colour change abilities (Tyrie et al., 2015, Smithers et al., 2018; Stevens & Ruxton, 2019; Eacock et al., 2019; Green et al., 2019). Being parts of an anti-predatory strategy, components of background matching are density-dependent, and are expected to evolve in response to variations of the demographic context, as during range expansions (McNamara and Houston, 1987; Brown & Kotler, 2004; Tollrian, 2015). Thus, analysing spatial patterns of inter-individual variation in background matching strategies within a well-characterized phylogeographic framework, might provide insights into the phenotypic consequences of past range dynamics.

In this study, we explore the effects of Pleistocene range dynamics on the evolution of anti-predatory traits, focusing on the tree frog *Hyla sarda*, an amphibian species using background matching to avoid predation. *H*.*sarda* is a tree frog endemic to the Tyrrhenian Islands (Western Mediterranean), whose evolutionary history was characterised by a major northward range expansion during the last glacial phase (Bisconti et al., 2011a, b; Spadavecchia et al., 2021). This tree frog colonised the northern portion of its current range (Corsica island and the Tuscan archipelago) from an ancestral area located along the central-eastern coast of Sardinia, taking advantage of a temporary land-bridge connection between Corsica and Sardinia (Spadavecchia et al., 2021). However, this land bridge was lost during the postglacial sea-level rise, preventing subsequent gene exchange between the source (Sardinian) and the newly colonised (Corsican) populations (Spadavecchia et al., 2021).

Here, we investigate whether Sardinian and Corsican populations of *H*.*sarda* differ in phenotypic traits linked to anti-predatory strategies. Specifically, we used common garden experiments to assess differences in the preference for alternative backgrounds (behavioural traits) and in the ability to change dorsal colour (physiological traits) among *H*.*sarda* populations.

## 2. Material and Methods

### Ethical note

Sampling procedures were performed under the approval of the Institute for Environmental Protection and Research ‘ISPRA’ (protocol # 5944), Ministry of Environment ‘MATTM’ (protocol #8275), Regione Sardegna (#12144) and Corsica (#2A20180206002 and #2B20180206001). Permission to temporarily house amphibians was granted by Local Health and Veterinary Centre, with license code 050VT427. All handling procedures outlined in the present study were approved by the Ethical Committee of the University of Tuscia for the use of live animals (D.R. n. 677/16 and D.R.644/17). All animals were released in the original sampling locations at the end of the experimentation.

### Sampling and housing

A total of 154 individuals were collected from 4 geographic areas, arranged along the inferred Late Pleistocene range expansion route of *H. sarda* (Spadavecchia et al., 2021; see Figure 1 and Table S1). Previous analyses of the bioclimatic niche of *H. sarda* carried out under both current and periglacial bioclimatic conditions, suggested spatially and temporally homogeneous habitat suitability models along the eastern coast (Bisconti et al., 2011b). Accordingly, to minimize this potential source of variation in our data (e.g. Stegen et al., 2004; Skold et al., 2013), all the individuals analysed were collected in strictly coastal sites (<10m above sea level; within 3 m from the coastline).

**Figure 1.**
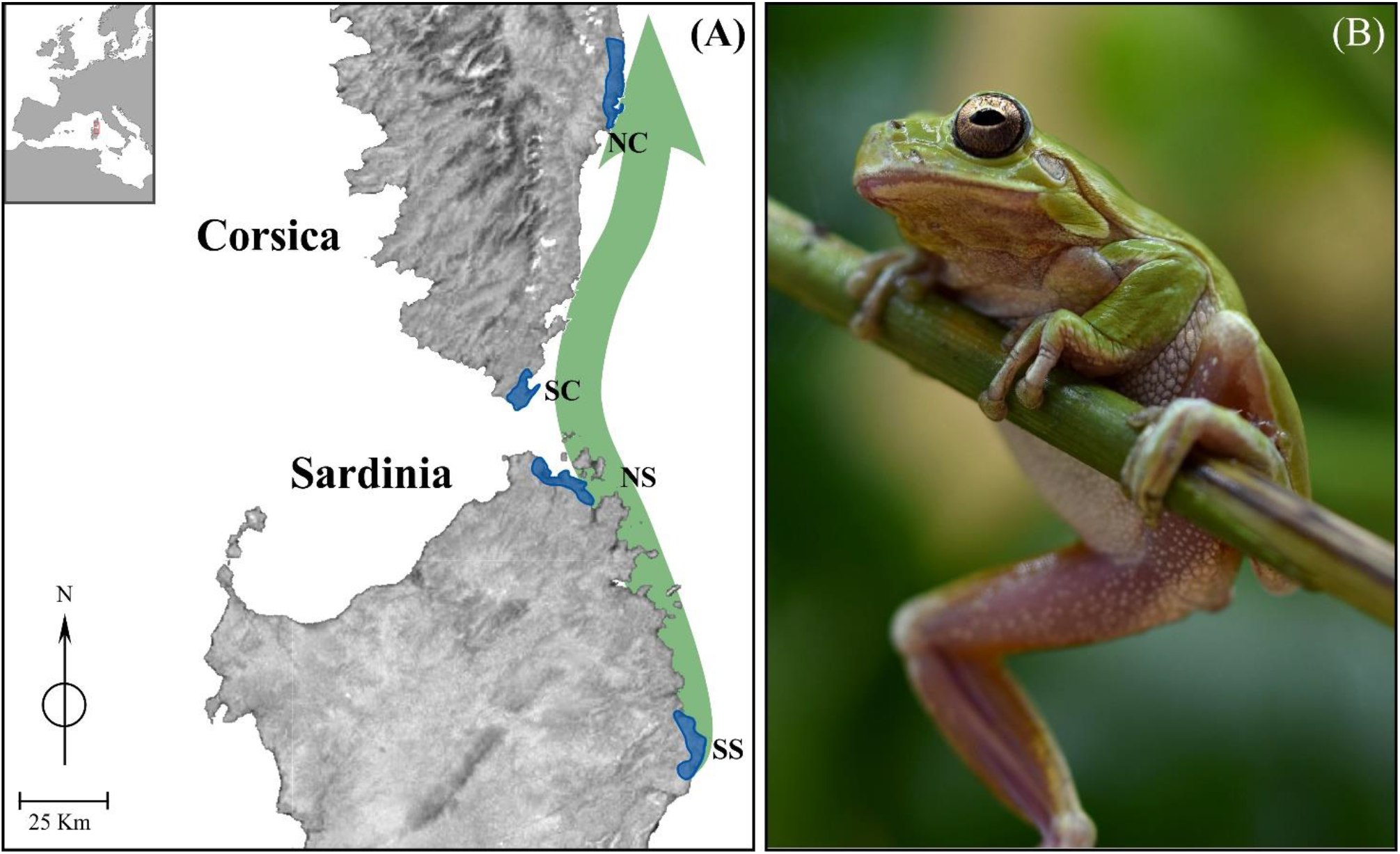
Study area and study species. **(A)** Blue shaded shapes indicate sampling areas; For more details about exact geographic locations, see TableS1. The green arrow shows the inferred Late Pleistocene range expansion route of *H. sarda*, according to Spadavecchia et al. (2021). **(B)** The Tyrrhenian tree frog *Hyla sarda*.

Individuals of *H. sarda* were transported to the laboratory within 3 days and then kept individually within cages (25 cm × 25 cm × 25 cm). All the tree frogs were provided with a water tank, and an oak wood as shelter, and were placed in a temperature-controlled room set to 24/25°C (monitored using a Hobo MX2201 temperature data-logger), 60%-80% of humidity, and natural photoperiod (Staniszewski, 1995). Individuals were fed ad libitum with crickets (Acheta domestica).

Before starting the tests, the animals were left undisturbed for two weeks except for routine feeding and cleaning duties. We also ascertained that individuals followed the normal day-night activity rhythms (higher activity at night-time); observations also included reactivity to food and calling activity at night. No adverse effects on the overall health of tree frogs or behavioural anomalies were observed during the experimental procedures.

For the assessment of individual differences in background matching strategies, two tests were performed: a colour change experiment, to measure individual ability to change colour (physiological trait), and a background choice experiment, to assess individual preference for alternative backgrounds (behavioural trait).

### Colour change experiments

To test individual abilities to change colour according to the background, each tree frog was exposed to two colour treatments, mimicking two extremes of the colour range encountered by tree frogs in their natural environment: 1) a brown treatment, and 2) a green treatment. Brown and green environments were obtained by positioning equally shaped brown and green plants, respectively, in the home cage (Réale et al., 2007). The acclimation period was spent in a colour-mixed environment, obtained by positioning an equal proportion of brown and green plants. During the first treatment, each tree frog was exposed to a brown environment, and the dorsal colouration of each individual was characterized; during the second treatment, tree frogs were exposed to a green environment, and dorsal colouration was characterized as well; finally, individual ability to change colour was quantified as the difference in colouration obtained after changing exposure from brown to the green background. Experiments were conducted inside the housing room, with a stable temperature to ensure that ambient temperature did not influence individual colour - since amphibians can change colour also for thermoregulation (Sköld et al., 2013). The exposure to each chromatic substrate lasted five days. After the exposure, the dorsal colouration of each individual was quantified using digital photography, a non-invasive method that prevents rapid colour change due to stress or handling (King et al.,1994, Sköld et al., 2013), and allows to post-process RAW format photographs (Stevens et al., 2007). To standardize image parameter quantification, individuals were positioned in a clear Petri dish fixed at 35 cm from the camera, close to a Datacolor SpyderChecker 24, for colour calibration, and were photographed using a Panasonic LUMIX (DMC-FZ300) digital camera.

Digital image analysis was performed using the plug-in Multispectral Image Calibration and Analysis Toolbox (Troscianko & Stevens, 2015) in IMAGEJ software (Schneider, 2012). For each photograph, we used white and black standards of known reflectance, respectively 90% and 4%, for normalisation. The normalisation of digital images is a procedure required to account for differences in illumination between photographs, and it is achieved by scaling each colour channel to a uniform reflectance level (Troscianko & Stevens., 2015). We sampled colour from the whole dorsum of each tree frog, by selecting a single region of interest (ROI), avoiding wrinkles and shadows, and measuring RGB (Süsstrunk et al., 1990; Hirsch, 2005) values of the ROI. As a measure of colour, we estimated the greenness (Eacock et al., 2017, 2019), calculated as the ratio between the cone catch values of the mediumwave and longwave photoreceptors [MW/(MW C LW)], which represent opponent mechanisms, following Arenas & Stevens (2017). From the RGB values, greenness was measured as G/(G+R)*100. Finally, individual ability to change colour was quantified as the difference in greenness values obtained after exposure to green and to brown background (hereafter Δ-greenness).

### Background choice experiment

Background choice and chromatic preferences were assessed by testing spontaneous behaviour. The experiment was carried out in the home cage (Réale et al., 2007), where individuals were provided with brown shelters and green plants. A visual scan sampling was conducted six times daily (9:00, 12:00, 15:00, 18:00, 21:00, 24:00) for one minute, for two days. The scan sampling covered the whole day because individual activity usually increases in tree frogs during the night. We estimated the percent time of inactivity (any discrete inactivity event longer than 3s measured as the proportion on total scans number) and the percent time of inactivity on green (hereafter percent time spent on a green substrate), defined as any discrete inactivity events on a green substrate longer than 3 s and measured as the proportion on the total number of inactivity events.

### Statistical analysis

Generalized linear models were carried out to assess the contribution of population geographic structure in explaining the observed variance in greenness values, in both brown and green environments, and in Δ-greenness. In each model, island (Sardinia vs Corsica) was included as fixed factor, and greenness as dependent variable. The analysis was run using the glm function in R package ‘nlme’ (Pinheiro et al., 2021, R version 3.5.3;), with a Gaussian distribution and with identity link function. For Δ-greenness, we used a Gamma distribution with an inverse function. We relied on the Akaike information criterion (AIC) to identify the distribution type and the link function improving model fitting.

We also applied generalized linear models to assess whether background choice significantly differed between the two islands (glm function in package ‘nlme’). Two different models were fitted, entering each variable singly (percent time of inactivity and percent time spent on a green substrate) and island as fixed factor. A Gaussian distribution with an identity link function was used for percent time of inactivity, while a Gamma distribution and a log link function were used for percent time spent on a green substrate.

## 3. Results

### Colour change experiment

We successfully collected photographs of the dorsal areas from 117 tree frog individuals. All the photographs were normalized and used for colour analysis. Over the entire dataset, greenness values spanned from 45.64 to 59.62, while Δ-greenness spanned from 0.00 to 8.01. The GLMs showed significant differences in colour change abilities between Sardinia and Corsica tree frogs (Table 1). After the exposure to the brown environment, greenness values were higher in Sardinian tree frogs (mean: 53.3, SE: ± 0.4) than in tree frogs from Corsica (mean: 49.9, SE: ± 0.4) (p-value < 0.001); see Fig 2A). The same pattern was found with the green substrate: individuals from Sardinia showed higher greenness values (mean: 51.8, SE: ± 0.2) than Corsican tree frogs (mean: 50.8, SE: ± 0.2) (p-value = 0.001; see Fig 2B). Finally, tree frogs from Sardinia showed higher Δ-greenness compared to the tree frogs from Corsica (mean: 2.47, SE: ± 0.26, and mean: 1.98, SE: ± 0.21, respectively), although these differences were not significant (p-value = 0.14).

**Table 1.** Outcome of generalized linear models using island as fixed factor.

| Variable | Coefficient | Standard Error | t-value | P-value |
| --- | --- | --- | --- | --- |
| Greenness on brown | 3.379 | 0.523 | 6.454 | **<0.0001** |
| Greenness on green | 0.922 | 0.282 | 3.269 | **0.0014** |
| Δ-greenness | -0.101 | 0.069 | -1.457 | 0.148 |
| % time of inactivity | -1.297 | 3.016 | -0.43 | 0.668 |
| % time spent on green | 0.327 | 0.146 | 2.237 | **0.028** |

**Figure 2.**
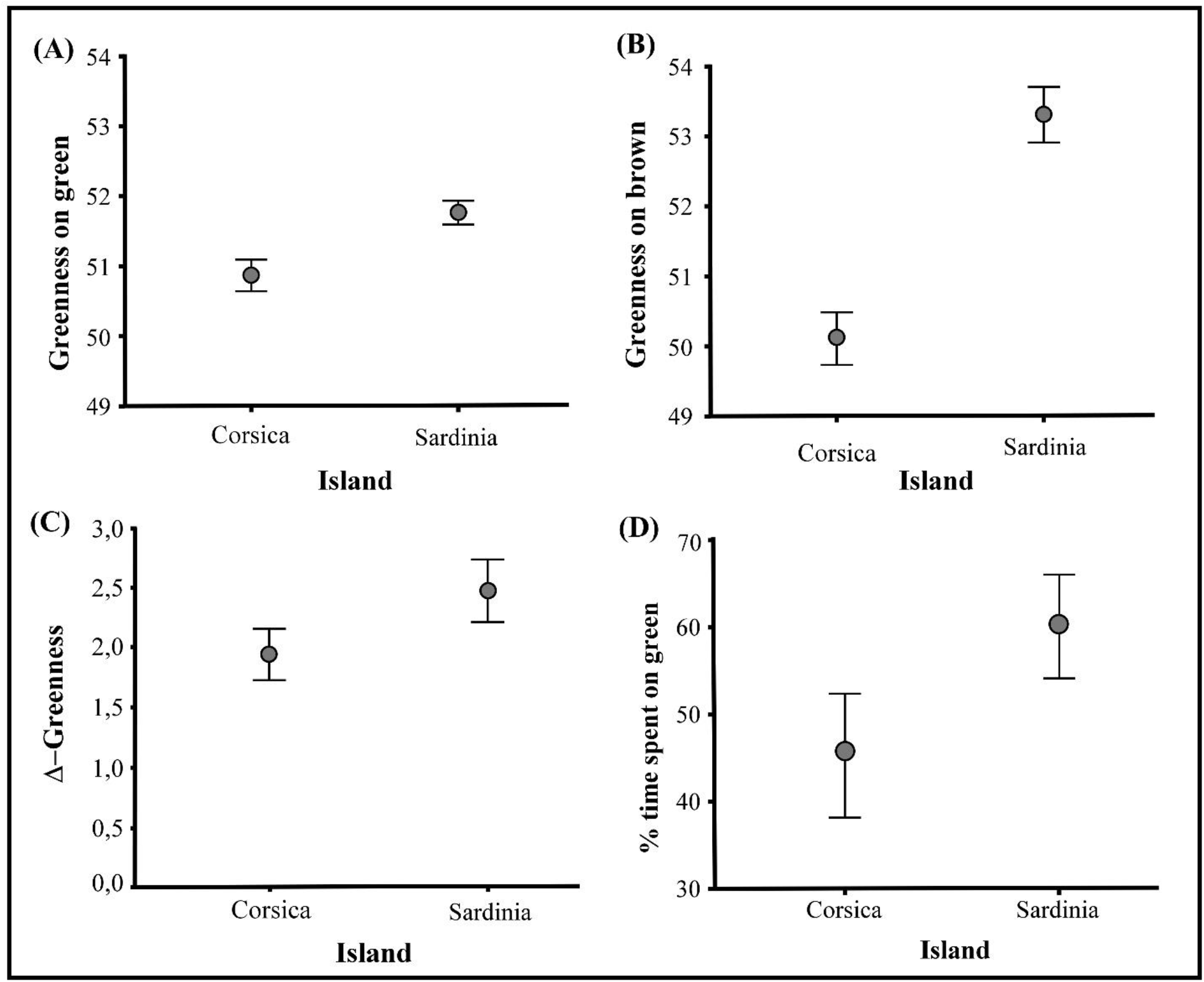
Geographic pattern of differentiation in greenness and in time spent on a green substrate. Estimated marginal means and standard errors obtained from generalized linear models for Corsican and Sardinian frogs. **(A)** Greenness values from the green test. **(B)** Greenness values from the brown test. **(C)** Δ-Greenness values indicating the differences between greenness values from the green test and brown test. **(D)** % time spent on green substrate.

### Background choice experiment

We extracted high quality data from 85 tree frogs individuals. Results from the background choice experiments revealed substantial inter-individual variation in the percent time spent on a green substrate. Overall, no significant differences in the percent time of inactivity were found between Sardinia and Corsica tree frogs (Table 1). On the contrary, GLMs indicated significant differences between tree frogs from the two islands in the percent time spent on the green substrate (Table 1). In particular, Sardinia tree frogs spent significantly more time on the green substrate compared to tree frogs from Corsica (Sardinia: mean: 4.07, SE: ± 0.10; Corsica: mean: 3.75, SE: ± 0.11; p-value = 0.028; see Fig. 2D).

## 4. Discussions

We explored the phenotypic consequences of a range expansion event, through the characterization of the geographic structure of anti-predatory traits along the Late Pleistocene range expansion route of *H. sarda*. We found significant differences in the ability to change dorsal colour and in background colour preferences between populations in Sardinia and Corsica. Although inter-individual and among-population differentiation in background matching abilities has been studied in other taxa (Marshall et al., 2015, 2016; Bailing et al., 2020; Yamamoto and Sota, 2020), including amphibians (Wente & Phillips, 2003; Kang et al., 2016; Barnett et al., 2021), this is the first study analysing patterns of variation in background matching abilities along a historical range expansion route. To the best of our knowledge, this is also the first study addressing spatial patterns of variation in anti-predatory traits in the context of the recent biogeographic history of populations.

Phenotypic differentiation among co-specific populations could be the outcome of non-adaptive processes, such as genetic drift associated with prolonged allopatric divergence or to founder events (Zhan et al., 2005; Amézquita, et al. 2009; Uyeda, 2009; Campbell et al., 2010). However, previous phylogeographic, historical demographic, and population genetic investigations (Bisconti et al., 2011a, 2011b; Spadavecchia et al., 2021), allow us to rule out these processes as major players in the case of the Tyrrhenian tree frog. Indeed, Corsican and Sardinian populations diverged less than 10k years ago, as suggested by isobaths and sea-level reconstructions (Spratt & Lisiecki, 2016). Accordingly, genetic investigations on both nuclear and mitochondrial markers did not show a remarkable genetic divergence between populations from Corsica and Sardinia (Bisconti et al., 2011a, 2011b). This pattern is expected in cases of a recent range expansion and when the time subsequent to the colonisation has not been sufficient for a new migration–drift equilibrium to be established (Hutchinson & Templeton, 1999). Likewise, the role of a founder effect can be excluded because the past land-bridge connecting the two main islands was wide and persistent throughout the last glacial phase (Thiede, 1978).

Anti-predatory traits might diverge among co-specific populations as a consequence of adaptive processes (Johnson & Belk, 2020). In the case of background matching, Sardinian and Corsican tree frogs might have diverged because of local adaptation to distinct chromatic environments. However, although the chromatic environment has not been characterised here, there are no prominent differences in habitat features and vegetation composition among Sardinian and Corsican tree frog habitats that could explain differential selective pressures (Ager et al., 2014). Furthermore, paleoclimatic reconstructions do not support a role for past habitat differences, as the coastland of the north of Sardinia (source population) and the south of Corsica (newly colonised) showed very similar bioclimatic conditions during the last glacial (Bisconti et al., 2011a).

On the other hand, the presence of marked phenotypic differences along a colonisation route suggests a role for past dispersal processes in shaping the spatial pattern of phenotypic variation in background matching. Indeed, range expansions represent a major challenge for organisms as they have to cope with new environmental conditions and, in turn, new eco-evolutionary constraints (Wilson et al., 2000; Sakai et al., 2001; Johnston & Temple, 2002; Steinhausen et al., 2008). Accordingly, density-dependent traits, like components of anti-predatory strategies, are expected to vary as a consequence of changes in the cost-benefit trade-off between safety and foraging, which depends on prey and predator densities (McNamara & Houston, 1987; Jeschke, 2006; Tollrian, 2015). During a range expansion, the ability to adjust camouflage is expected to be a critical factor in determining the success of colonisations (Lodge, 1993; Schoener & Spiller, 1995), as individuals with higher proficiency in background matching should have a higher chance to survive and to establish the new populations (Fairchild & Howell, 2004; Duarte et al., 2017). However, the proficiency in background matching could be limited because of physiological costs and energetic constraints associated with colour change, and the stress of dealing with a new environment (Hill & McGraw, 2006; Polo-Cavia & Gomez-Mestre, 2017; Moreno-Rueda, 2020). Noteworthy, it has been recently suggested that when phenotypic plasticity is costly, selection can actively eliminate it canalizing towards the favoured phenotype (Levis & Pfennig, 2019). Thus, the selection pressures at the front of the expansion wave, coupled with modified life history trade-offs in the transient demographic conditions occurring at the expansion front, might have shaped the observed pattern of phenotypic differentiation. This scenario is in accordance with results from studies conducted on invasive species, which support the hypothesis that dispersal influences personality, morphology, performance, physiology (Gruber et al., 2017; Cameron et al., 2019; Hudson et al., 2020; Kosmala et al., 2020) but also anti-predator response (Hudson et al., 2017).

Interestingly, recent studies on geographic variation in traits affecting physiology and ageing in *H. sarda* showed similar spatial patterns of variation (Liparoto, 2020; Canestrelli et al., 2021). This spatial congruence suggests the intriguing possibility that the range expansion event left traces on multiple phenotypic traits, thus promoting the evolution of a dispersal syndrome (Ronce & Clobert, 2012; Stevens et al., 2012, 2013).

In conclusion, our study provides new insights on the influence of a Late Pleistocene range expansion in shaping the spatial pattern of phenotypic variation in the Tyrrhenian tree frog. These results support the hypothesis that past range expansions (and other phylogeographic processes) might have played a prominent, albeit largely underexplored role in shaping the evolution of phenotypes and their spatial patterns of differentiation among intraspecific populations (Bonte et al., 2012; Clobert et al., 2012; Zamudio et al., 2016; Canestrelli et al., 2016a, b).

## Acknowledgements

We thank Alessandro Carlini, Giacomo Grignani, Lorenzo Latini, Armando Macali, for assistance with sampling and the experimental procedures. This work was supported by Italian Ministry of Education, University and Research (PRIN project 2017KLZ3MA).

## Authors contributions

D.Can. conceived the study. GS, AL, RB collected the data. GS, AC, D.Can., D.Cos. and RB, performed formal analysis and interpreted the results. GS wrote the original draft of the manuscript. AC, RB and D.Can. reviewed the manuscript. All authors approved the final manuscript.

## References

Ager, A. A., Preisler, H. K., Arca, B., Spano, D., & Salis, M. (2014). Wildfire risk estimation in the Mediterranean area. Environmetrics, 25(6), 384–396.

Amezquita, A., Lima, A. P., Jehle, R., Castellanos, L., Ramos, O., Crawford, A. J., & Hoedl, W. (2009). Calls, colours, shape, and genes: a multi-trait approach to the study of geographic variation in the Amazonian frog Allobates femoralis. Biological journal of the linnean society, 98(4), 826–838.

Arenas, L. M., & Stevens, M. (2017). Diversity in warning coloration is easily recognized by avian predators. Journal of evolutionary biology, 30(7), 1288–1302.

Avise, J. C. (2009). Phylogeography: retrospect and prospect. J. Biogeography. 36, 3–15.

Baling, M., Stuart-Fox, D., Brunton, D. H., & Dale, J. (2020). Spatial and temporal variation in prey color patterns for background matching across a continuous heterogeneous environment. Ecology and Evolution, 10(5), 2310–2319

Barnett, J. B., Michalis, C., Scott-Samuel, N. E., & Cuthill, I. C. (2021). Colour pattern variation forms local background matching camouflage in a leaf-mimicking toad. Journal of Evolutionary Biology, 34(10), 1531–1540.

Bates, D., Mächler, M., Bolker, B., Walker, S. (2015). Fitting Linear Mixed-Effects Models Using lme4. Journal of Statistical Software, 67(1), 1–48.

Bisconti, R., Canestrelli, D., & Nascetti, G. (2011b). Genetic diversity and evolutionary history of the Tyrrhenian treefrog Hyla sarda (Anura: Hylidae): adding pieces to the puzzle of Corsica– Sardinia biota. Biological Journal of the Linnean Society, 103(1), 159–167.

Bisconti, R., Canestrelli, D., Colangelo, P., & Nascetti, G. (2011a). Multiple lines of evidence for demographic and range expansion of a temperate species (Hyla sarda) during the last glaciation. Molecular Ecology, 20(24), 5313–5327.

Bonte, D., Van Dyck, H., Bullock, J. M., Coulon, A., Delgado, M., Gibbs, M., & Travis, J. M. (2012). Costs of dispersal. Biological reviews, 87(2), 290–312.

Brown, J. S., & Kotler, B. P. (2004). Hazardous duty pay and the foraging cost of predation. Ecology letters, 7(10), 999–1014.

Cameron, M. S., & Donald, J. A. (2019). Different vasodilator mechanisms in intermediate-and small-sized arteries from the hindlimb vasculature of the toad Rhinella marina. American Journal of Physiology-Regulatory, Integrative and Comparative Physiology, 317(3), R379–R385.

Campbell, P., Pasch, B., Pino, J. L., Crino, O. L., Phillips, M., & Phelps, S. M. (2010). Geographic variation in the songs of neotropical singing mice: testing the relative importance of drift and local adaptation. Evolution: International Journal of Organic Evolution, 64(7), 1955–1972.

Canestrelli, D., Bisconti, R., & Carere, C. (2016b). Bolder takes all? The behavioral dimension of biogeography. Trends in ecology & evolution, 31(1), 35–43.

Canestrelli, D., Bisconti, R., Liparoto, A., Angelier, F., Ribout, C., Carere, C., & Costantini, D. (2021). Biogeography of telomere dynamics in a vertebrate. Ecography.

Canestrelli, D., Porretta, D., Lowe, W. H., Bisconti, R., Carere, C., & Nascetti, G. (2016a). The tangled evolutionary legacies of range expansion and hybridization. Trends in Ecology & Evolution, 31(9), 677–688.

Chiocchio, A., Arntzen, J. W., Martínez-Solano, I., de Vries, W., Bisconti, R., Pezzarossa, A., & Canestrelli, D. (2021). Reconstructing hotspots of genetic diversity from glacial refugia and subsequent dispersal in Italian common toads (Bufo bufo). Scientific Reports, 11(1), 1–14.

Clobert, J., Baguette, M., Benton, T. G., and Bullock, J. M. (2012). Dispersal ecology and evolution. Oxford: Oxford University Press.

Cobben, M. M. P., Verboom, J., Opdam, P. F. M., Hoekstra, R. F., Jochem, R., and Smulders, M. J. M. (2015). Spatial sorting and range shifts: consequences for evolutionary potential and genetic signature of a dispersal trait. Journal of Theoretical Biology, 373, 92–99.

Duarte, R. C., Flores, A. A., & Stevens, M. (2017). Camouflage through colour change: mechanisms, adaptive value and ecological significance. Philosophical Transactions of the Royal Society B: Biological Sciences, 372(1724), 20160342.

Eacock, A., Rowland, H. M., Edmonds, N., & Saccheri, I. J. (2017). Colour change of twig-mimicking peppered moth larvae is a continuous reaction norm that increases camouflage against avian predators. PeerJ, 5, e3999.

Eacock, A., Rowland, H. M., van’t Hof, A. E., Yung, C. J., Edmonds, N., & Saccheri, I. J. (2019). Adaptive colour change and background choice behaviour in peppered moth caterpillars is mediated by extraocular photoreception. Communications biology, 2(1), 1–8.

Fairchild, E. A., & Howell, W. H. (2004). Factors affecting the post-release survival of cultured juvenile Pseudopleuronectes americanus. Journal of Fish Biology, 65, 69–87.

Fløjgaard, C., Normand, S., Skov, F., & Svenning, J. C. (2009). Ice age distributions of European small mammals: insights from species distribution modelling. Journal of Biogeography, 36(6), 1152–1163.

Green, S. D., Duarte, R. C., Kellett, E., Alagaratnam, N., & Stevens, M. (2019). Colour change and behavioural choice facilitate chameleon prawn camouflage against different seaweed backgrounds. Communications biology, 2(1), 1–10.

Gruber, J., Brown, G., Whiting, M. J., & Shine, R. (2017). Geographic divergence in dispersal-related behaviour in cane toads from range-front versus range-core populations in Australia. Behavioral Ecology and Sociobiology, 71(2), 38.

Hewitt, G. M. (1996). Some genetic consequences of ice ages, and their role in divergence and speciation. Biological journal of the Linnean Society, 58(3), 247–276.

Hewitt, G. M. (2000). The genetic legacy of the Quaternary ice ages. Nature, 405(6789), 907–913.

Hewitt, G. M. (2004a). Genetic consequences of climatic oscillations in the Quaternary. Philosophical Transactions of the Royal Society of London. Series B: Biological Sciences, 359(1442), 183–195.

Hewitt, G. M. (2011a). Mediterranean peninsulas: the evolution of hotspots. In Biodiversity hotspots (pp. 123–147). Springer, Berlin, Heidelberg.

Hewitt, G. M. (2011b). Quaternary phylogeography: the roots of hybrid zones. Genetica, 139(5), 617–638.

Hill, G. E., McGraw, K. J., 2006. Bird coloration. Vol. I: Mechanisms and measurements. Harvard University Press, Cambridge.

Hirsch, R. (2004). Exploring colour photography: a complete guide. Laurence King Publishing.

Hofreiter, M., & Stewart, J. (2009). Ecological change, range fluctuations and population dynamics during the Pleistocene. Current biology, 19(14), R584–R594.

Hudson, C. M., Brown, G. P., & Shine, R. (2017). Evolutionary shifts in anti-predator responses of invasive cane toads (Rhinella marina). Behavioral Ecology and Sociobiology, 71(9), 1–9.

Hudson, C. M., Vidal-García, M., Murray, T. G., & Shine, R. (2020). The accelerating anuran: evolution of locomotor performance in cane toads (Rhinella marina, Bufonidae) at an invasion front. Proceedings of the Royal Society B, 287(1938), 20201964.

Hutchison, D. W., & Templeton, A. R. (1999). Correlation of pairwise genetic and geographic distance measures: inferring the relative influences of gene flow and drift on the distribution of genetic variability. Evolution, 53(6), 1898–1914.

Jeschke, J. M. (2006). Density-dependent effects of prey defenses and predator offenses. Journal of theoretical biology, 242(4), 900–907.

Johnson, J. B., & Belk, M. C. (2020). Predators as agents of selection and diversification. Diversity, 12(11), 415.

Johnston, I. A., & Temple, G. K. (2002). Thermal plasticity of skeletal muscle phenotype in ectothermic vertebrates and its significance for locomotory behaviour. Journal of Experimental Biology, 205(15), 2305–2322.

King, R.B., Hauff, S. & Phillips, J. B. (1994). Physiological color change in the green treefrog: responses to background brightness and temperature. Copeia 1994, 422–432.

Kjernsmo, K., & Merilaita, S. (2012). Background choice as an anti-predator strategy: the roles of background matching and visual complexity in the habitat choice of the least killifish. Proceedings of the Royal Society B: Biological Sciences, 279(1745), 4192–4198.

Kosmala, G. K., Brown, G. P., & Shine, R. (2020). Thin-skinned invaders: geographic variation in the structure of the skin among populations of cane toads (Rhinella marina). Biological Journal of the Linnean Society, 131(3), 611–621.

Levis, N. A., & Pfennig, D. W. (2019). Phenotypic plasticity, canalization, and the origins of novelty: evidence and mechanisms from amphibians. In Seminars in cell & developmental biology (Vol. 88, pp. 80–90). Academic Press.

Liparoto, A., Canestrelli, D., Bisconti, R., Carere, C., & Costantini, D. (2020). Biogeographic history moulds population differentiation in ageing of oxidative status in an amphibian. Journal of Experimental Biology, 223(21).

Lodge, D. M. (1993). Biological invasions: lessons for ecology. Trends in ecology & evolution, 8(4), 133–137.

Lucati, F., Poignet, M., Miró, A., Trochet, A., Aubret, F., Barthe, L., & Ventura, M. (2020). Multiple glacial refugia and contemporary dispersal shape the genetic structure of an endemic amphibian from the Pyrenees. Molecular Ecology, 29(15), 2904–2921.

Marshall, K. L., Philpot, K. E., & Stevens, M. (2016). Microhabitat choice in island lizards enhances camouflage against avian predators. Scientific Reports, 6(1), 1–10.

Marshall, K. L., Philpot, K. E., Damas-Moreira, I., & Stevens, M. (2015). Intraspecific colour variation among lizards in distinct island environments enhances local camouflage. PLoS One, 10(9), e0135241.

McNamara, J. M., & Houston, A. I. (1987). Starvation and predation as factors limiting population size. Ecology, 68(5), 1515–1519.

Merilaita, S., & Stevens, M. (2011). Crypsis through background matching. Animal camouflage: mechanisms and function, 17-33.

Merilaita, S., Scott-Samuel, N. E., & Cuthill, I. C. (2017). How camouflage works. Philosophical Transactions of the Royal Society B: Biological Sciences, 372(1724), 20160341.

Miller, T. E., Angert, A. L., Brown, C. D., Lee-Yaw, J. A., Lewis, M., Lutscher, F., & Williams, J. L. (2020). Eco-evolutionary dynamics of range expansion. Ecology, 101(10), e03139.

Moreno-Rueda, G. (2020). The evolution of crypsis when pigmentation is physiologically costly.

Pinheiro, J. Bates, D., DebRoy, S. Sarkar, D., R Core Team (2021). nlme: Linear and Nonlinear Mixed Effects Models. R package version 3.1-153, https://CRAN.R-project.org/package=nlme.

Polo-Cavia, N., & Gomez-Mestre, I. (2017). Pigmentation plasticity enhances crypsis in larval newts: associated metabolic cost and background choice behaviour. Scientific reports, 7(1), 1–10.

Réale, D., Reader, S. M., Sol, D., McDougall, P. T., and Dingemanse, N. J. (2007). Integrating animal temperament within ecology and evolution. Biological reviews, 82 (2), 291–318.

Ronce, O., & Clobert, J. (2012). Dispersal syndromes. Dispersal ecology and evolution, 155, 119–138.

Ruxton, G. D., Allen, W. L., Sherratt, T. N., & Speed, M. P. (2019). Avoiding attack: the evolutionary ecology of crypsis, aposematism, and mimicry. Oxford University Press.

Sakai, A. K., Allendorf, F. W., Holt, J. S., Lodge, D. M., Molofsky, J., With, K. A., & Weller, S. G. (2001). The population biology of invasive species. Annual review of ecology and systematics, 32(1), 305–332.

Schmitt, T. (2007). Molecular biogeography of Europe: Pleistocene cycles and postglacial trends. Frontiers in zoology, 4(1), 1–13.

Schneider, C.A., Rasband, W.S., Eliceiri, K.W. (2012). NIH Image to ImageJ: 25 years of image analysis. Nature Methods 9, 671–675.

Schoener, T. W., & Spiller, D. A. (1995). Effect of predators and area on invasion: an experiment with island spiders. Science, 267(5205), 1811–1813.

Sköld, H. N., Aspengren, S. & Wallin, M. (2013). Rapid color change in fish and amphibians– function, regulation, and emerging applications. Pigment Cell Melanoma Res. 26, 29–38.

Smithers, S. P., Rooney, R., Wilson, A., & Stevens, M. (2018). Rock pool fish use a combination of colour change and substrate choice to improve camouflage. Animal Behaviour, 144, 53–65.

Spadavecchia, G., Chiocchio, A., Bisconti, R., & Canestrelli, D. (2021). Paso doble: a two-step Late Pleistocene range expansion in the Tyrrhenian tree frog Hyla sarda. Gene, 780, 145489.

Spratt, R. M., & Lisiecki, L. E. (2016). A Late Pleistocene sea level stack. Climate of the Past, 12(4), 1079–1092.

Staniszewski, M. (1995). Amphibians in captivity. TFH.

Stegen, J. C., Gienger, C. M., & Sun, L. (2004). The control of color change in the Pacific tree frog, Hyla regilla. Canadian Journal of Zoology, 82(6), 889–896.

Steinhausen, M. F., Sandblom, E., Eliason, E. J., Verhille, C., & Farrell, A. P. (2008). The effect of acute temperature increases on the cardiorespiratory performance of resting and swimming sockeye salmon (Oncorhynchus nerka). Journal of Experimental Biology, 211(24), 3915–3926.

Stevens, M. (2016). Color change, phenotypic plasticity, and camouflage. Frontiers in Ecology and Evolution, 4, 51.

Stevens, M., & Ruxton, G. D. (2019). The key role of behaviour in animal camouflage. Biological Reviews, 94(1), 116–134.

Stevens, M., Párraga, C. A., Cuthill, I. C., Partridge, J. C., & Troscianko, T. S. (2007). Using digital photography to study animal coloration. Biological Journal of the Linnean society, 90(2), 211–237.

Stevens, M., Troscianko, J., Wilson-Aggarwal, J. K., & Spottiswoode, C. N. (2017). Improvement of individual camouflage through background choice in ground-nesting birds. Nature ecology & evolution, 1(9), 1325–1333.

Stevens, V. M., Trochet, A., Blanchet, S., Moulherat, S., Clobert, J., & Baguette, M. (2013). Dispersal syndromes and the use of life-histories to predict dispersal. Evolutionary applications, 6(4), 630–642.

Stevens, V. M., Trochet, A., Van Dyck, H., Clobert, J., & Baguette, M. (2012). How is dispersal integrated in life histories: a quantitative analysis using butterflies. Ecology letters, 15(1), 74–86.

Stuart-Fox, D., & Moussalli, A. (2011). 13 Camouflage in colour-changing animals. Animal camouflage: mechanisms and function, 237.

Süsstrunk, S., Buckley, R., & Swen, S. (1999). Standard RGB color spaces. In Color and Imaging Conference (Vol. 1999, No. 1, pp. 127–134). Society for Imaging Science and Technology.

Szűcs, M., Vahsen, M. L., Melbourne, B. A., Hoover, C., Weiss-Lehman, C., & Hufbauer, R. A. (2017). Rapid adaptive evolution in novel environments acts as an architect of population range expansion. Proceedings of the National Academy of Sciences, 114(51), 13501–13506.

Taberlet, P., Fumagalli, L., Wust-Saucy, A. G., & Cosson, J. F. (1998). Comparative phylogeography and postglacial colonization routes in Europe. Molecular ecology, 7(4), 453–464.

Thiede, J. (1978). A glacial Mediterranean. Nature, 276(5689), 680–683.

Tollrian, R., Duggen, S., Weiss, L. C., Laforsch, C., & Kopp, M. (2015). Density-dependent adjustment of inducible defenses. Scientific Reports, 5(1), 1–9.

Troscianko, J., & Stevens, M. (2015). Image calibration and analysis toolbox–a free software suite for objectively measuring reflectance, colour and pattern. Methods in Ecology and Evolution, 6(11), 1320–1331.

Tyrie, E. K., Hanlon, R. T., Siemann, L. A., & Uyarra, M. C. (2015). Coral reef flounders, Bothus lunatus, choose substrates on which they can achieve camouflage with their limited body pattern repertoire. Biological Journal of the Linnean Society, 114(3), 629–638.

Uyeda, J. C., Arnold, S. J., Hohenlohe, P. A., & Mead, L. S. (2009). Drift promotes speciation by sexual selection. Evolution: International Journal of Organic Evolution, 63(3), 583–594.

Weiss, S., & Ferrand, N. (2007). Phylogeography of southern European refugia (pp. 341–357). Dordrecht: springer.

Wilson, R. S., James, R. S., & Johnston, I. A. (2000). Thermal acclimation of locomotor performance in tadpoles and adults of the aquatic frog Xenopus laevis. Journal of Comparative Physiology B, 170(2), 117–124.

Yamamoto, N., & Sota, T. (2020). Evolutionary fine-tuning of background-matching camouflage among geographical populations in the sandy beach tiger beetle. Proceedings of the Royal Society B, 287(1941), 20202315.

Zamudio, K. R., Bell, R. C., & Mason, N. A. (2016). Phenotypes in phylogeography: Species’ traits, environmental variation, and vertebrate diversification. Proceedings of the National Academy of Sciences, 113(29), 8041–8048.

Zhan, J., Linde, C. C., Jürgens, T., Merz, U., Steinebrunner, F., & McDonald, B. A. (2005). Variation for neutral markers is correlated with variation for quantitative traits in the plant pathogenic fungus Mycosphaerella graminicola. Molecular Ecology, 14(9), 2683–2693.

