## Supplementary material for "Spatial differentiation of background matching strategies along a Late Pleistocene range expansion route": Table S1

**Table S1.** Geographic coordinates of sampling locations and sample size for individuals of *Hyla sarda* analysed in this study.

| Island | Area | Latitude N | Longitude E | N |
|---|---|---|---|---|
| Sardinia | SS | 40° 26′ | 09° 47′ | 27 |
| | NS | 41° 11′ | 09°19′ | 46 |
| Corsica | SC | 41° 26′ | 09°12′ | 40 |
| | NC | 42° 16′ | 09°31′ | 41 |
